# Small immunological clocks identified by Deep Learning and Gradient Boosting

**DOI:** 10.1101/2022.10.28.514283

**Authors:** Alena Kalyakulina, Igor Yusipov, Elena Kondakova, Maria Giulia Bacalini, Claudio Franceschi, Maria Vedunova, Mikhail Ivanchenko

## Abstract

**Background:** The aging process affects all systems of the human body, and the observed increase in inflammatory components affecting the immune system in old age can lead to the development of age-associated diseases and systemic inflammation.

**Results:** We propose a small clock model SImAge based on a limited number of immunological biomarkers. To regress the chronological age from cytokine data, we first use a baseline Elastic Net model, gradient-boosted decision trees models, and several deep neural network architectures. For the full dataset of 46 immunological parameters, DANet, SAINT, FT-Transformer and TabNet models showed the best results for the test dataset. Dimensionality reduction of these models with SHAP values revealed the 10 most age-associated immunological parameters, taken to construct the SImAge small immunological clock. The best result of the SImAge model shown by the FT-Transformer deep neural network model has mean absolute error of 6.94 years and Pearson *ρ* = 0.939 on the independent test dataset. Explainable artificial intelligence methods allow for explaining the model solution for each individual participant.

**Conclusions:** We developed an approach to construct a model of immunological age based on just 10 immunological parameters, coined SImAge, for which the FT-Transformer deep neural network model had proved to be the best choice. The model shows competitive results compared to the published studies on immunological profiles, and takes a smaller number of features as an input. Neural network architectures outperformed gradient-boosted decision trees, and can be recommended in the further analysis of immunological profiles.

## 1 Introduction

### 1.1. Background

The immune system plays an important role in protecting the human organism from various infections. An increase in inflammatory components affecting the innate and adaptive arms of the immune system [1–4] is observed in old age, which may be associated with the development of various age-associated diseases [5–8] and systemic inflammation [9–14]. This chronic inflammation, caused by the metabolic products of damaged cells and environmental influences, has been associated with aging and leads to progressive damage of many tissues [5, 15]. This phenomenon of increased levels of circulating inflammatory mediators is called inflammaging [16–18]. However, there are no standard biomarkers to fully characterize it [19, 20].

The aging process is extremely complex and involves all body systems, so there have been many attempts to characterize it and estimate its rate from different perspectives: combining different clinical parameters, predicting the risk of mortality or cognitive deterioration [21–25]. A broad class of aging biomarkers are various chronological age predictors, or clocks. Epigenetic clocks, which attempt to predict a person’s age based on DNA methylation data [26–29], are the most common. Recently, inflammatory clocks have also been developed [30–32]. Both genetic factors involved in the formation of immune composition [33–35] and environmental factors [36, 37] can lead to significant individual differences in immune characteristics. An individual’s immune status continuously changes over time; therefore, immune age can vary even among healthy individuals of the same chronological age [38, 39]. Understanding the relationship between biological aging and an individual’s immunological profile and predisposition to age-associated diseases may facilitate the development of tools to slow aging and improve longevity [40].

One of the first immunological age biomarkers is Inflammatory Biologic Age, which uses 9 inflammatory markers to construct an aggregate age estimate based on data from over 3,000 subjects using the Klemera-Doubal method [21]. This inflammatory age acceleration has been shown to be associated with increased risks of cardiovascular disease and mortality. The original work compares mean values with standard deviations of clinical and inflammatory ages; no error values are given, so an explicit comparison with other approaches is difficult. Another aggregated biomarker of immunological aging is IMM-AGE, which reflects multidimensional changes in immune status with age and is significantly associated with mortality. The association of IMM-AGE score with cardiovascular disease has also been shown [30]. Aggregating information on cellular phenotyping and cytokine responses in a fairly wide age range of individuals (40-90 years), IMM-AGE demonstrates a P value of 10^-60 for linear regression with age. However, this score does not explicitly estimate age. Another recently developed metric for human immunological age is iAge, an inflammatory aging clock [31]. It uses information on cytokines, chemokines and growth factors from a thousand of healthy and diseased subjects ranging from age 8 to 96 years to produce an aggregated age estimate using a guided autoencoder. This nonlinear method, a type of deep neural network, eliminates the noise and redundancy of the raw data while preserving important biological information. Nevertheless, the model still requires the entire input data to be fed to the input, and therefore is not entirely compact. Despite a quite considerable average error of 15.2 years, the model manifests associations with multimorbidity, immune aging, frailty, and cardiovascular aging. Another example is the ipAGE model, which is sensitive to age-associated acceleration in chronic kidney disease [32]. Constructed using the classic Elastic Net approach and relying on 38 immunological biomarkers, ipAGE demonstrates a mean absolute error of 7.27 years and root mean squared error of 8.33 years for the test subset of the control group and detects immunological biomarkers significantly differentially expressed between cases and controls. Notably, the model better assesses the phenotype of accelerated aging in patients with end-stage renal disease by quantifying inflammaging than various epigenetic clock models. It, however, has limitations due to the small training sample size and the lack of cross-validation. Another class of aggregated immunological biomarkers constructs age estimates based on peptides or plasma proteins data [41, 42]. Such models are characterized by a large dimensionality of input data, ranging from hundreds or thousands of human plasma proteins to several million peptide features, that may be difficult to assess in clinical applications.

All the described immunological age predictors use different quantitative measures, and the input data for all of the models are tabular: the rows contain samples (individual participants) and the columns contain features (immunological measures). One of the most common methods for constructing age predictor models (clocks) for different types of data is Elastic Net. This model is used as the basis for multiple age predictor models based on epigenetic [26–29], immunological [32, 42], transcriptomic [43], metabolomic [44–46], microRNA [47], and proteomic [48] data. The more advanced methods like gradient boosting are also actively used to construct age predictors, particularly from epigenetic [49] or gut microbiome [50] data. In the last few years, deep learning methods, including neural networks, autoencoders, and other approaches are also actively used to construct various clock model, to name epigenetic [51], immunological [31], transcriptomic [52], gut microbiome [53] and hematological [54–56].

Linear models, like Elastic Net, are easily interpretable and widely used in age prediction tasks. Despite this, they have a number of drawbacks: assume linear relationships between dependent and independent variables, do not take into account possible nonlinear relationships, and are sensitive to outliers [57]. In this regard, making use of the gradient-boosted decision trees (GBDT) and neural network approaches has considerable potential, in particular for tabular data. Among GBDTs the most frequent choice for classification and regression problems for tabular data is XGBoost [58], LightGBM [59] and CatBoost [60], cf. also [61, 62]. Several neural network architectures have been developed specifically for tabular data, to compete with GBDT models. The increasing number of heterogeneous data estimation tasks with both continuous and categorical features, for which the traditional approaches are not always well applied, calls for flexible solutions, in particular based on deep neural networks that can capture complex nonlinear dependencies in the data [63]. Regularization-based approaches [64, 65], different transformer architectures [66–68], and hybrid methods that combine the advantages of tree and neural network models [69] have been developed to handle tabular data. Despite the many approaches that have been proposed, there is still no consensus on which methods perform best for tabular datasets [61–63, 68].

Simple linear models are easy to interpret, the importance of individual features is determined by the values of the corresponding coefficients; most of the treelike models are also interpretable. For neural network models the situation is more complicated: not all existing models allow to obtain the importance ranking of individual attributes. This can be mitigated by explainable artificial intelligence (XAI) approaches that can determine the contribution of individual features to the final prediction for almost any model [70–76]. Explainability can also be used to reduce the dimensionality of models. Discarding unimportant, noisy features can often reduce the input dimensionality of a model and improve the results. Global and local types of explainability are especially interesting [77–79]. Global explainability helps to interpret the behavior of the model as a whole: which features have the highest impact on the prediction of certain classes (for classification) or specific values (for regression). Local explainability helps to determine why the model made its prediction for a particular sample, and how this was influenced by the feature values for that sample. This type of model explainability meets the need of personalized medicine.

### 1.2. Study design and novelty

The primary goal of this work is to develop a small clock based on a limited number of immunological biomarkers. We use the classic Elastic Net model as the baseline, as the most common method for constructing age predictors on various types of biological data. To explore more advanced tools for chronological age regression on tabular data we make use of gradient-boosted decision trees (GBDT), which stand among the most well-proven approaches specifically for tabular data. In many problems GBDT models showed similar or better results in comparison with deep learning models, but required less tuning and showed better computational performance [61]. Nevertheless, we also considered several neural network architectures designed to solve problems on tabular data.

Linear models, in particular Elastic Net, are easy to interpret, whereas tree and neural network models are highly nonlinear, much more complex, and require methods of explainable artificial intelligence for interpretability. Accordingly, we first apply XAI methods to determine the most important features for age prediction models based on immunological data. We also use the obtained ranking to build portable models that consist of a small number of immunological biomarkers. As a rationale, they are much easier to implement with regard to practical applications, particularly, in regard to cost. Above that, XAI methods have the potential of explaining the model decision for each particular sample (person), meeting the challenges of personalized medicine.

## 2. Results

### 2.1. Data description

#### Participants

The study that involved a control group of 300 healthy volunteers recruited in the Nizhny Novgorod region in 2019-2021 (train/validation dataset, 260 samples) and 2022-2023 (test dataset, 40 samples) was performed at Lobachevsky State University of Nizhny Novgorod. Exclusion criteria were fairly loose, limiting chronic diseases in the acute phase, oncological diseases, acute respiratory viral infections, and pregnancy only. The sex distribution has a significant predominance of women: 173 women and 87 men are in the train and validation datasets, 27 women and 13 men in the test dataset. The age distribution was also not uniform, with an overall age range from 19 to 101 years, participants from the age group of 30-60 years prevailed (Figure 1(A)).

**Figure 1.**
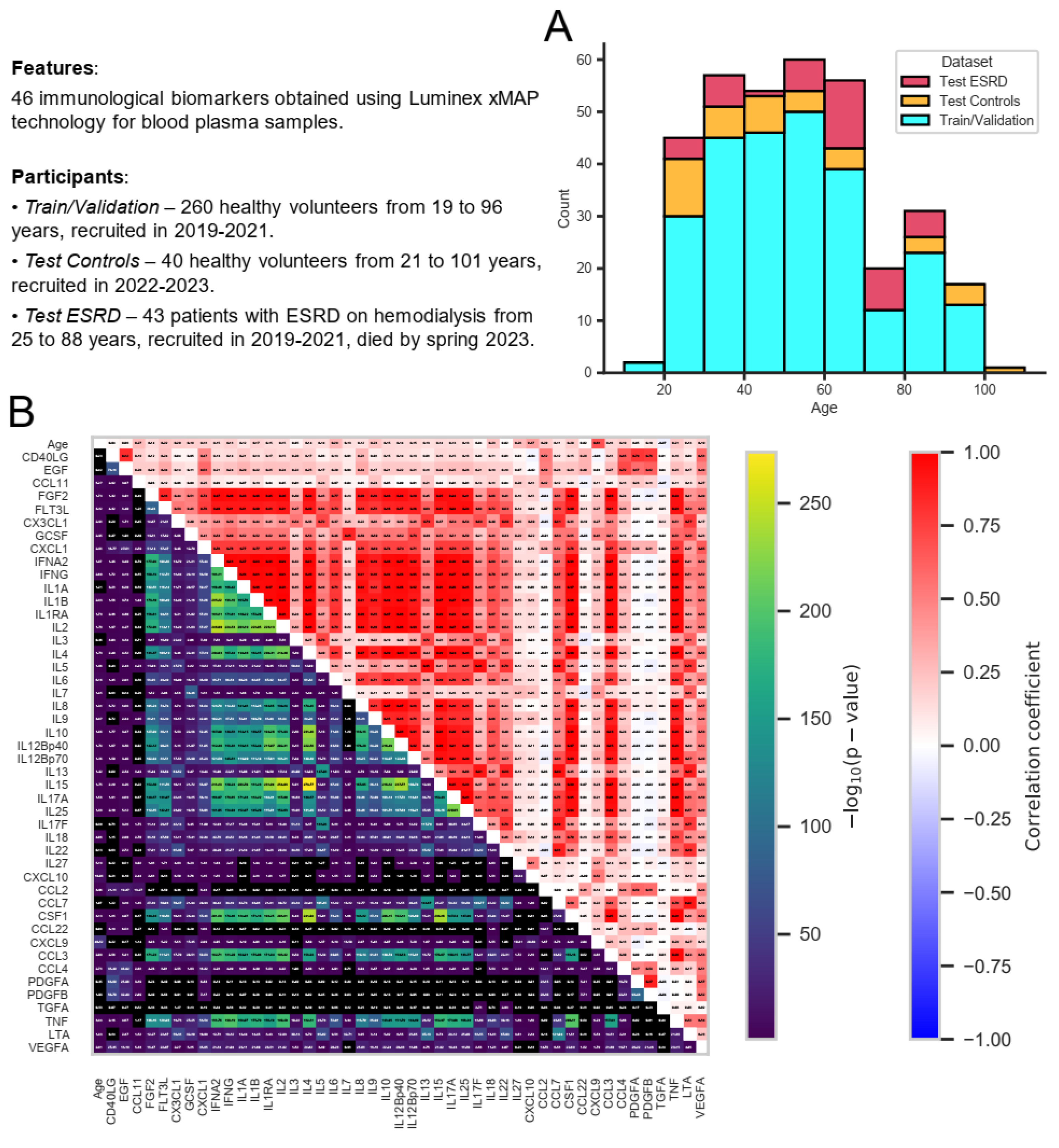
Information about the data analyzed in this study. (A) Stacked histogram showing distribution of samples by age and group: train/validation, test control, test ESRD. (B) Relationships between all immunological biomarkers and age for all control participants (from both train/validation and test datasets): Pearson correlation coefficient (top triangle) and the p-value for testing non-correlation (bottom triangle) were calculated for each pair of features. All p-values were adjusted using Benjamini-Hochberg FDR-correction (p-value<0.05).

Separate test dataset included 43 ESRD (end-stage renal chronic disease) patients on hemodialysis recruited in 2019-2021 at “FESPHARM NN” hemodialysis centers in the Nizhny Novgorod region, Russia [32], who did not survive until May 2023 because of the effects of the underlying disease.

#### Features

Figure 1(B) displays 46 immunological biomarkers considered in this study for all control samples (both train/validation and test datasets). They were obtained using Luminex xMAP technology for blood plasma samples (see Methods Section 4.1 for details). To examine the possible correlations between these immunological parameters and their correlation with age, Pearson correlation coefficients and p-values for testing non-correlation (Benjamini-Hochberg FDR-corrected p-values<0.05) were calculated for each pair of features. CXCL9 has proved to be the most correlated with age, in agreement with the earlier studies [31, 32, 80, 81]. Recent evidence suggests that CXCL9 is an important marker of inflammation and plays a key role in the development of age-associated diseases, such as neurodegeneration [82], chronic kidney disease [32], glaucoma [83] and various inflammatory diseases [84–86]. In addition, there emerged a whole group of interleukin biomarkers that are significantly correlated with each other (the most strongly correlated pairs are IL2 vs IL17A, IL2 vs IL15, IL17A vs IL15 with Pearson correlation coefficients almost equal to one). The relationship between IL2 and IL15 has been demonstrated [87], including cancer therapy [88, 89], as well as the relationship between IL2 and IL17A [90, 91], and between IL15 and IL17A [92].

All data used in the present work, such as immunological biomarker values, age, status (healthy control or ESRD) and sex of participants, are summarized in Supplementary Table 1.

### 2.2. Experiment design

The aim of this study is to develop a machine learning model that solves the problem of regression of chronological age (in other words, clocks) on a reduced set of the most significant immunological biomarkers (Small Immuno Age - SImAge). Initially, there are 46 immunological markers available, and the goal is to effectively reduce their number in such a way so that the accuracy of the age predictor model does not decrease significantly. On the implementation side, a number of manufacturers (i.e. Merck KGaA, Bio-Rad) are able to produce custom panels with a pre-selected set of biomarkers (for example, specific 10 immunological parameters instead of 46), that can significantly reduce its cost.

The developed workflow is shown in Figure 2. The first step is to train the models using all available immunological biomarkers (Figure 2, step 1). We used a variety of machine learning models, ranging from classical linear regression with Elastic Net regularization (a popular choice in age regression tasks [26, 27, 32, 43, 45, 48]) to GBDT and various DNN (deep neural network) architectures specifically designed for tabular data. The GBDT and DNN models employed in this paper are the state-of-the-art models that compete for better results in many studies [61, 63, 67, 68]. One of the goals of this study is to find out which class of models is the best in our specific problem setting with a limited number of samples. The detailed description of the used models is given in Section 4.2. 5-fold cross-validation and hyperparametric search were used to determine the model with optimal parameters that provided the lowest mean absolute error (MAE) on the validation dataset (Section 4.3). After that, the best model (with lowest validation MAE) was tested on the separate test dataset. The main metric for model comparison is the MAE on this test dataset, upon which the final ranking of the models is based. The value of the Pearson correlation coefficient was also tracked along with the MAE both during cross-validation with hyperparametric search and during testing the model on the independent dataset. This allows us to avoid situations, when a small error is achieved only for the most representative age range.

**Figure 2.**
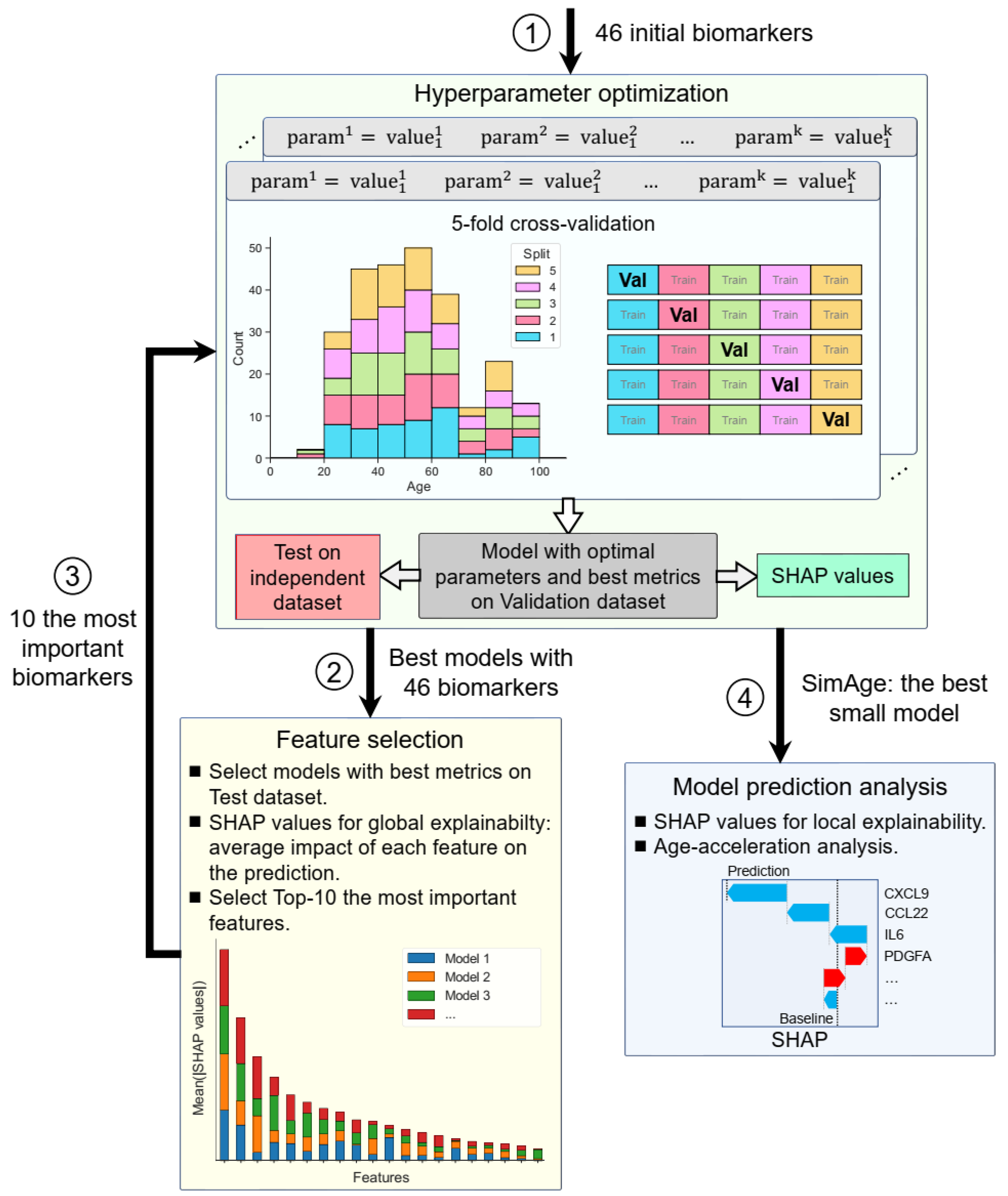
The main steps of the performed analysis. Step 1: Building baseline machine learning models on a complete list of 46 immunological biomarkers. For each type of models 5-fold cross-validation and hyperparametric search are used to determine the best model with optimal parameters in terms of MAE and Pearson correlation coefficient on validation dataset. Then, these best models are tested on the independent test dataset and the final model ranking is built according to MAE on the test dataset. The actual splits are shown in the histogram. Step 2: Identification of the most important features from the best baseline models. Averaged absolute SHAP values for the best models, calculated for train/validation datasets together, are used to select the most important immunological biomarkers for building small models. Step 3: Building small models on a short list of selected immunological biomarkers. As for Step 1, 5-fold cross-validation with hyperparametric search is used for determining the best model on validation dataset. SImAge is the best model on the selected biomarkers in terms of MAE on the test dataset. Step 4: Analysis of age predictions produced by SImAge. The contribution of individual immunological parameter values to individual participant predictions (using local explainability and SHAP values) is measured, as well as the importance of features in groups of people with age acceleration (immunological age obtained by SImAge model is higher than chronological age) and age deceleration (SImAge is lower than chronological age).

After several top baseline models were identified, we selected the most important features (Figure 2, step 2) according to their importance in global explainability described by SHAP values (Section 4.4). For this purpose, the best models with the lowest MAE for the test dataset were first identified. Then the features were ranked by their averaged absolute SHAP values, calculated for train/validation datasets together, in these best models, yielding top 10 features in result.

To build the best portable model for predicting human immunological age (SImAge), we used the same types of models that emerged at the previous step. However, the models took only the 10 best parameters determined earlier as an input. 5-fold cross validation with hyperparametric search was also performed to find the best small model (Section 4.3). As for the first step (but for 10 features, not 46) the best model was tested on the independent dataset, and Pearson correlation coefficient was tracked with MAE. As a result, we chose a model with the best MAE on the test dataset, and coined it SImAge (Figure 2, step 3).

Then, we investigated the predictions produced by the SImAge model (Figure 2, step 4). Using SHAP values (Section 4.4), we estimated the contribution of certain immunological biomarker values to the model predictions for individual participants. The cumulative contribution of the individual features to the predictions with positive and negative age acceleration was determined.

### 2.3. Baseline results

We implemented the models from three conceptually different classes to solve the problem of regression of chronological age from immunological profile data: (i) the classical linear Elastic Net model, (ii) Gradient Boosted Decision Trees: XGBoost [58], LightGBM [59], CatBoost [60] and (iii) Deep Neural Networks: simple Multilayer Perceptron (MLP), Neural Additive Models (NAM) [93], Neural Oblivious Decision Ensembles (NODE) [69], Deep Abstract Networks (DANet) [94], TabNet [65], Automatic Feature Interaction Learning via Self-Attentive Neural Networks (AutoInt) [66], Self-Attention and Intersample Attention Transformer (SAINT) [67], Feature Tokenizer and Transformer (FT-Transformer) [68]. A detailed description of these models is given in Section 4.2.

For each model, an experiment with a particular configuration of model parameters included 5 cross-validation splits (details in Methods, Section 4.3). Within each split, an individual model was trained on the training dataset (80%) and validated on the validation dataset (20%). Based on 5 splits, the mean and standard deviation of MAE and Pearson correlation coefficient on the validation dataset were calculated. This experiment was repeated many times with different combinations of model parameters during hyperparametric search, whose main aim is to determine the optimal combination of parameters with the minimal MAE (more details in the methods, Section 4.3). Besides this, the minimal MAE and corresponding Pearson correlation coefficient values for the best split was saved. Then the models were tested on the independent test dataset and the ranking of the models were built according to the result of this testing. This step of the analysis is represented schematically within the overall study design in Figure 2, step 1.

Table 1 shows the results of solving the age prediction problem for all baseline models trained on the full set of 46 immunological biomarkers. Despite that GBDT models and neural network architectures show similar results on the validation dataset, on the test dataset DNNs have a significant advantage - DANet, SAINT, FT-Transformer and TabNet models show the highest results (MAE on test dataset less than 8 years), with a large gap between them and all other considered models. GBDT models on test dataset show MAE of more than 10 years, which may indicate their weaker ability to generalize compared to DNNs. It is interesting to note that three out of four top models implement an attention mechanism (SAINT, FT-Transformer, and TabNet). In all cases, the results of the best models significantly outperform the classical linear regression model with ElasticNet regularization, usually implemented in age prediction problems.

**Table 1.** Results of age prediction by baseline models using all 46 immunological biomarkers. For each model, the average MAE and Pearson correlation coefficient *ρ* with corresponding standard deviations, as well as the best MAE and *ρ* for the validation dataset are given. For each model, the best MAE and *ρ* for the test dataset are given. The best models with the lowest MAE values on the test dataset are highlighted. Angular brackets represent average values.

| Type | Model | Test MAE | Test $\rho$ | Validation $\langle \text{MAE} \rangle \pm \text{STD}$ | Validation $\langle \rho \rangle \pm \text{STD}$ | Validation Best MAE | Validation Best $\rho$ |
| --- | --- | --- | --- | --- | --- | --- | --- |
| Linear | Elastic Net | 15.09 | 0.607 | $12.95 \pm 1.28$ | $0.584 \pm 0.194$ | 11.71 | 0.738 |
| GBDT | XGBoost | 10.85 | 0.881 | $8.97 \pm 0.56$ | $0.838 \pm 0.022$ | 8.22 | 0.859 |
| | LightGBM | 11.44 | 0.886 | $8.16 \pm 0.98$ | $0.857 \pm 0.033$ | 6.81 | 0.904 |
| | CatBoost | 10.93 | 0.893 | $8.93 \pm 1.70$ | $0.831 \pm 0.062$ | 6.94 | 0.904 |
| DNN | MLP | 10.77 | 0.825 | $8.78 \pm 0.84$ | $0.838 \pm 0.016$ | 7.55 | 0.856 |
| | NAM | 9.57 | 0.897 | $9.78 \pm 0.98$ | $0.802 \pm 0.053$ | 8.12 | 0.873 |
| | NODE | 10.96 | 0.879 | $10.09 \pm 1.30$ | $0.773 \pm 0.048$ | 8.67 | 0.841 |
| | **DANet** | **7.16** | **0.956** | $8.86 \pm 1.73$ | $0.813 \pm 0.053$ | 6.63 | 0.904 |
| | **TabNet** | **7.91** | **0.914** | $8.68 \pm 0.90$ | $0.825 \pm 0.036$ | 7.46 | 0.884 |
| | AutoInt | 12.65 | 0.698 | $10.59 \pm 1.03$ | $0.760 \pm 0.046$ | 9.33 | 0.789 |
| | **SAINT** | **7.24** | **0.942** | $9.73 \pm 1.16$ | $0.792 \pm 0.059$ | 7.82 | 0.867 |
| | **FT-Transformer** | **7.45** | **0.937** | $9.65 \pm 1.29$ | $0.806 \pm 0.047$ | 7.49 | 0.880 |

### 2.4. Feature selection and dimensionality reduction

Linear machine learning models, like Elastic Net, are easy to interpret: the greater the absolute value of the feature’s weight coefficient, the more important this feature is. The considered GBDT models have a built-in functionality to determine the feature importance, unlike most of the considered neural network architectures (only NAM and TabNet architectures have such built-in functionality). The same model-agnostic approach, based on the calculation of SHAP (SHapley Additive exPlanations) values, is used to obtain the feature importance in all the analyzed models [95]. This is a game theoretic approach to explaining the results of any machine learning model, which links the optimal credit allocation to local explanations using classical Shapley values from game theory and related extensions (details in Methods, Section 4.4) [96].

SHAP values can show how each individual feature affects the final prediction of the model (age estimation in our case), positively or negatively. Using this approach, the feature importance values (corresponding to mean absolute SHAP values, calculated for train/validation datasets together) were calculated for 4 best models (DANet, SAINT, FT-Transformer, TabNet). Figure 3(A) shows the stacked histogram for all immunological features and their feature importance values. As a result, the most robust 10 immunological parameters with the highest summarized averaged absolute SHAP values were selected. These biomarkers are then used to construct a portable immunological clock. This step of the study is shown schematically as a part of the general flowchart in Figure 2, step 2. In the literature, the following biomarkers from the Top-10 IL6 [97, 98], CSF1 [99], PDGFA [100], CXCL10 [101] were associated with aging, while the rest CXCL9 [31], CCL22 [102], PDGFB [103], VEGFA [104] were associated with different age-related and inflammatory pathological conditions.

**Figure 3.**
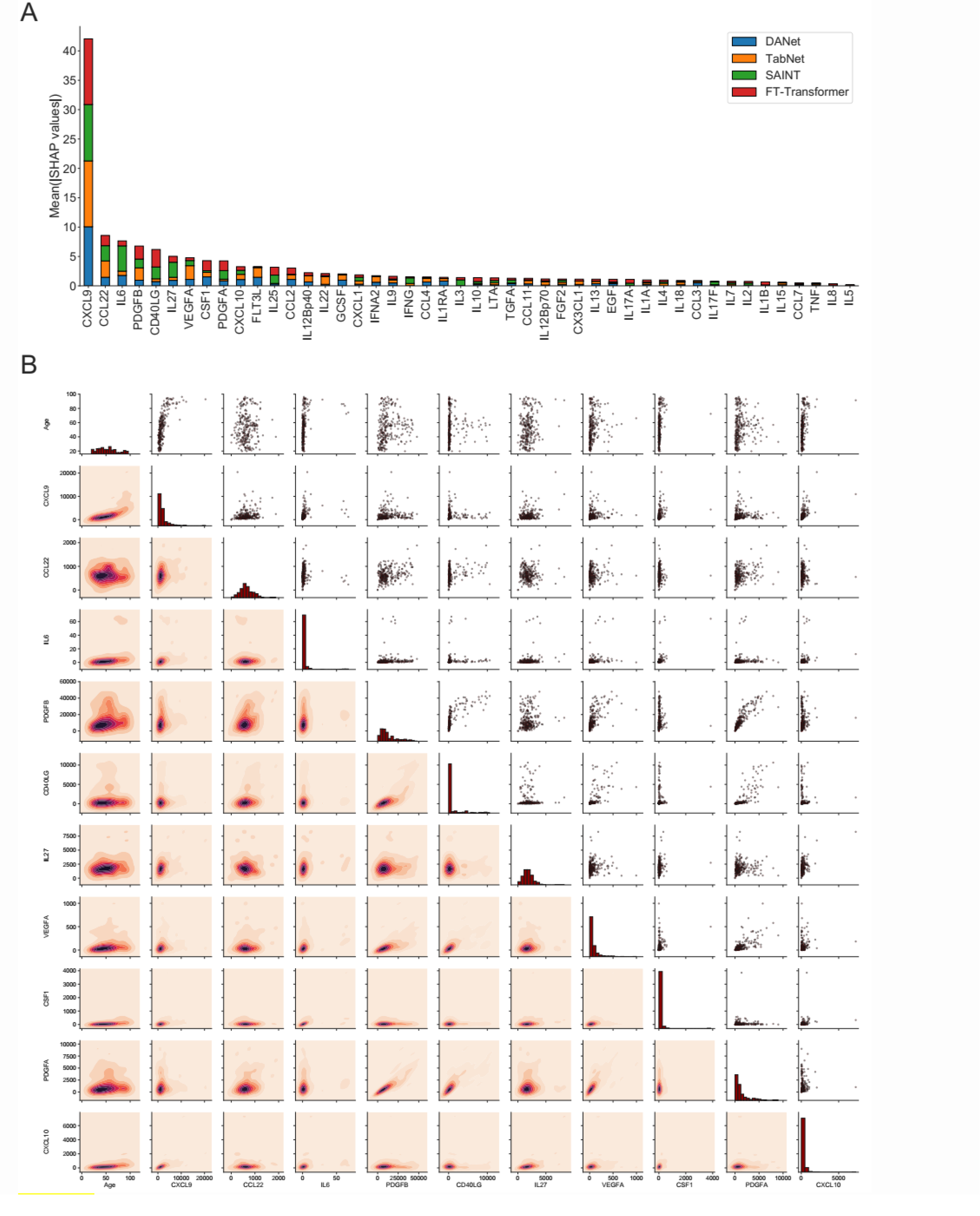
The 10 most important immunological features that were selected for the construction of the small immunological clocks. (A) Ranking of the features according to their averaged absolute SHAP values in the best models: DANet (blue), TabNet (orange), SAINT (green), FT-Transformer (red). The 10 selected biomarkers with the highest importance values are taken for building small models. (B) Relative distribution of 10 most important immunological biomarker values and chronological age. The diagonal elements illustrate the distribution of each individual feature, the scatter plots in the upper triangle and the probability density functions in the lower triangle illustrate the relationships between each pair of features.

Figure 3(B) shows the relative distribution of the 10 selected immunological biomarker values and chronological age. The diagonal elements illustrate the distribution of each individual feature, the scatter plots in the upper triangle and the probability density functions in the lower triangle illustrate the relationship between each pair of features. Expectantly, there is a clear proportional relationship between PDGFA and PDGFB, since both biomarkers belong to the group of platelet-derived growth factor (PDGF), growth factors for fibroblasts, smooth muscle cells and glial cells [105]. The values of the Pearson correlation coefficients and the corresponding p-values for these biomarkers can be found in Figure 1(B). It should also be noted that the biomarkers from the above-mentioned set of mutually correlated interleukins are not found among the 10 selected features.

### 2.5. Small immunological clocks (SImAge)

Portable chronological age regression models are constructed for the selected 10 most significant immunological features. We restrict our attention to the 4 models that demonstrated the best baseline results on the test dataset: DANet, SAINT, FT-Transformer, TabNet.

Like in the baseline experiments, for each model we performed 5-fold cross-validation with hyperparametric search to find the best models with optimal parameters giving the lowest MAE on the validation dataset. Next, all best models were tested on the independent dataset and the SImAge model was determined as the model with the lowest MAE on the test dataset using 10 immunological biomarkers. This step of the pipeline is reflected in the general flowchart in Figure 2, step 3.

Table 2 shows the results for the DANet, SAINT, FT-Transformer and TabNet models, which received 10 selected biomarkers as input. The best result was obtained for the FT-Transformer model as highlighted in the table. It should be noted that the other models demonstrated reasonably close performance. Comparing the results to those obtained on the full set of parameters, we conclude that the selected 10 immunological parameters can serve a solid basis to obtain a valid age estimation.

**Table 2.** Results of age prediction by models based on 10 selected immunological biomarkers. For each model, the average MAE and Pearson correlation coefficient (*ρ*) with corresponding standard deviations, as well as the best MAE and *ρ* for the validation dataset are given. For each model, the best MAE and *ρ* for the test dataset are given. The best model with the lowest MAE value on the test dataset is highlighted. Angular brackets represent average values.

| Model | Test MAE | Test $\rho$ | Validation $\langle \text{MAE} \rangle \pm \text{STD}$ | Validation $\langle \rho \rangle \pm \text{STD}$ | Validation Best MAE | Validation Best $\rho$ |
| --- | --- | --- | --- | --- | --- | --- |
| DANet | 7.19 | 0.942 | $8.92 \pm 1.00$ | $0.830 \pm 0.034$ | 7.36 | 0.881 |
| TabNet | 8.08 | 0.910 | $8.86 \pm 1.30$ | $0.830 \pm 0.042$ | 7.67 | 0.873 |
| SAINT | 8.05 | 0.912 | $8.97 \pm 1.41$ | $0.825 \pm 0.049$ | 8.21 | 0.863 |
| **FT-Transformer** | **6.94** | **0.939** | $8.74 \pm 1.05$ | $0.831 \pm 0.033$ | 7.21 | 0.888 |

The best model predicting age on a small number of immunological biomarkers SImAge (FT-Transformer) has a MAE=6.94 years and Pearson correlation coefficient 0.939 on the test dataset. Parameters of this model are presented in Supplementary Table 2. Figure 4(A) shows the predicted versus chronological age for all datasets: train, validation, test controls, test ESRD. It can be seen that samples from the datasets of controls (train, validation, test controls) are located along the bisector of SImAge=Age, while ESRD samples have significantly higher SImAge prediction than their chronological age. Figure 4(B) illustrates the distribution of SImAge acceleration in train, validation, test controls and test ESRD datasets, which is defined as the residual relative to linear approximation for all controls datasets. It can be seen that ESRD group has a significant SImAge acceleration (p-value=4.58e-08) in relation to the control group.

**Figure 4.**
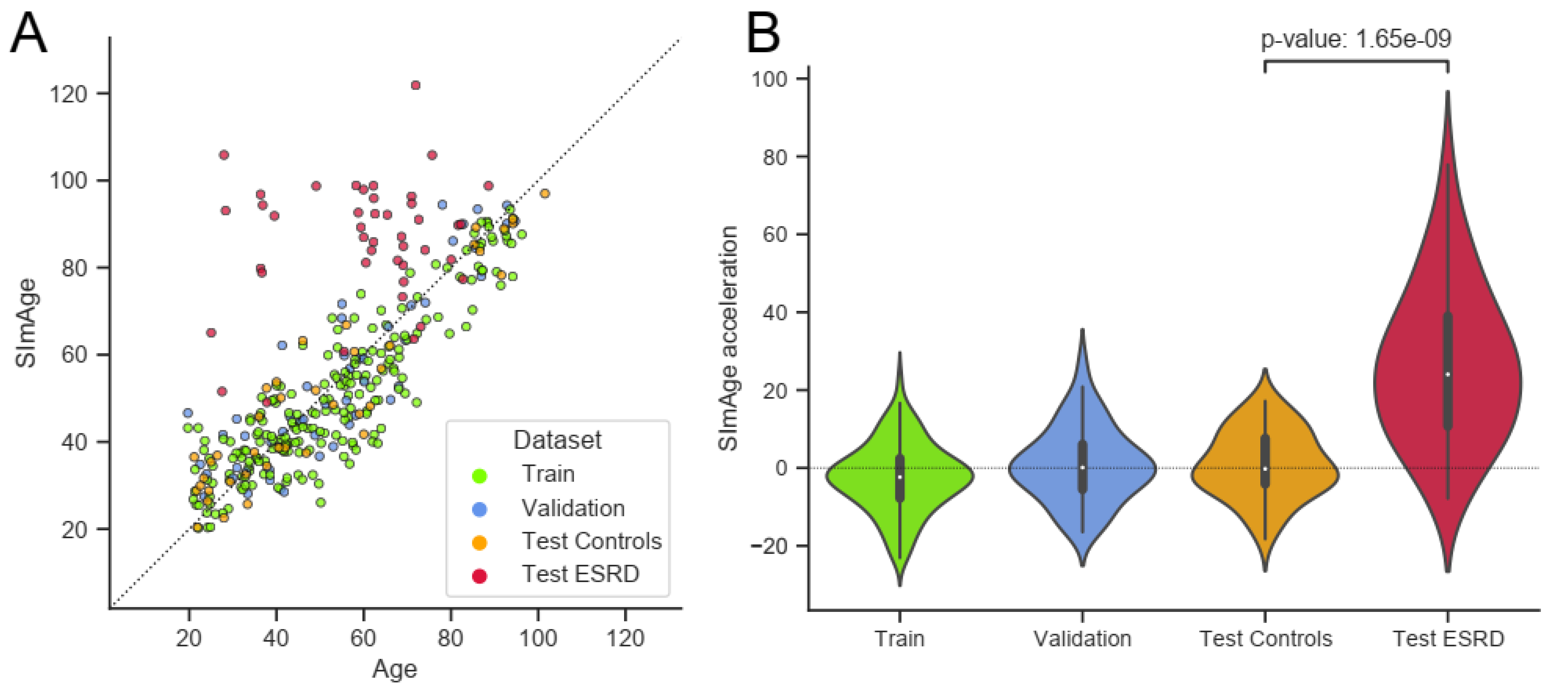
Results for the best model predicting age on a small number of immunological biomarkers (SImAge). (A) Scatter plot representing result of SImAge prediction for train (green), validation (blue), test controls (orange) and test ESRD (red) datasets versus chronological age. The black solid line is the bisector of SImAge=Age. (B) Violin plots showing distribution of SImAge acceleration in train (green), validation (blue), test controls (orange) and test ESRD (red) datasets. SImAge acceleration is calculated as residuals relative to linear approximation for all controls together (train, validation, test controls).

We also analyzed MAE of the SImAge model for different age ranges, taking into account all control participants (from train, validation and test datasets). The results were slightly different between the groups, but no age-related dependency of the MAE value was observed. Significant differences between males and females were not also found. The detailed results are given in Supplementary Table 3.

The test dataset with ESRD patients, who died from the effects of the underlying disease, was considered to address the sensitivity of the proposed small immunological clock to mortality. As shown above (Figure 4), there is a significant positive age acceleration in this group, it persists in all age ranges and is present in both sexes (Supplementary Table 3). This can be evidence that significant positive acceleration for age estimates of the proposed clocks may be a sign of higher mortality risks. Further investigations are required to validate its association to the other cause of mortality.

### 2.6. Model predictions analysis

In addition to characterizing feature importance, SHAP values can also be used to highlight local explainability, increasing the transparency of a particular prediction of the model. SHAP values also help to explain the result for a given sample, and to determine the contribution of individual feature values to the prediction.

SHAP values show how a particular value of a selected feature for a certain sample changes the basic prediction of the model (the average prediction of the model in the background dataset, which is train and validation datasets together in our case). This approach is most clearly illustrated with waterfall plots, which show summation of SHAP values towards an individual prediction (Figure 5). It manifests which features affected the change in model predictions relative to the mean value in each case, and characterizes the cumulative positive and negative contributions of all the features (which can lead to a rather large model error). Figure 5(A) shows an example case of the correct model prediction for a control sample with a small age, while Figure 5(B) shows it for a control sample with a large age. In both cases we can see that the most important and age-correlated biomarker CXCL9 significantly shifts the model prediction (small values shift negatively, large values shift positively), with PDGFA(B) also in the top, and CCL22 significant for a young person, IL6 contributing for the older one. Figure 5(C, D) shows control examples of model predictions with large negative and large positive age accelerations, respectively. Interestingly, in these particular cases, the highest influence on the age mismatch was caused by CD40LG value, which is only fifth by importance in the ranking (see Figure 3A). Figure 5(E, F) shows ESRD examples of model predictions with small and large positive age accelerations, respectively. CXCL9 has the highest influence on both predictions with IL6 and CSF1 standing next in the line.

**Figure 5.**
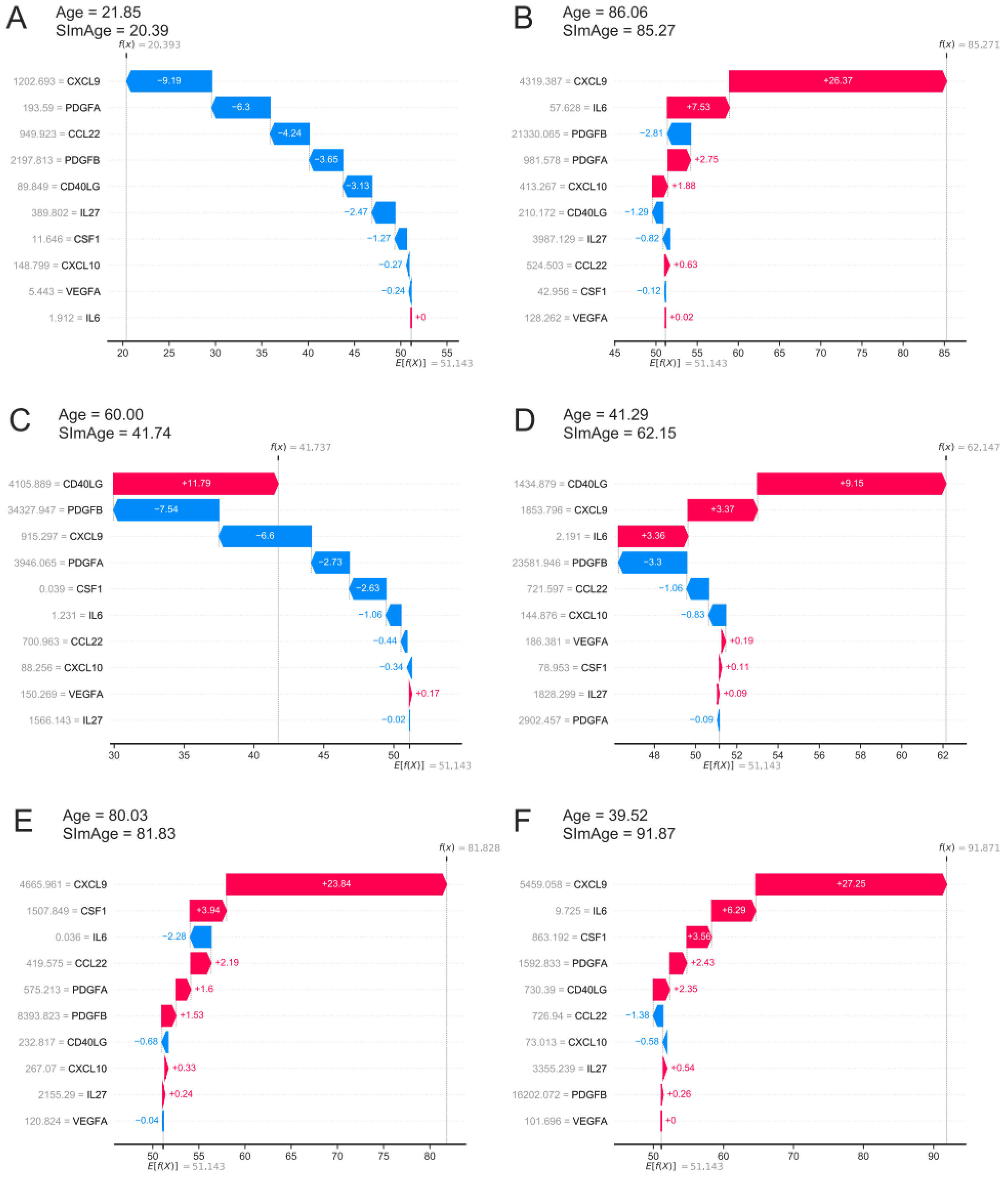
The local explainability of the SImAge model based on SHAP values is illustrated by waterfall plots. The bottom part of each waterfall plot starts with the expected value of the model output *E*[*f*(*X*)] (the average prediction of this model on the background dataset). Each row shows by how much in the positive (red) or negative (blue) direction each feature shifts the prediction relative to the expected value to the final model prediction for that sample *f*(*X*). (A) Example of a control sample with low age acceleration and young age. (B) Example of a control sample with low age acceleration and old age. (C) Example of a control sample with a high negative age acceleration. (D) Example of a control sample with high positive age acceleration. (E) Example of an ESRD sample with low age acceleration. (F) Example of an ESRD sample with high positive age acceleration.

Further on, we investigated the cumulative effect of individual features in predicting significant positive and negative age acceleration. All control participants were divided into 3 groups with different types of age acceleration (acceleration was calculated as the difference between SImAge and chronological age: Acceleration = SImAge – Age): with absolute value of age acceleration less than MAE (weak acceleration), with value of age acceleration less than -MAE (significant negative acceleration), with value of age acceleration greater than MAE (significant positive acceleration). ESRD participants were also analyzed (Figure 6(A)). For each group, the total contribution of individual immunological parameters to the final prediction was analyzed (the average value of absolute SHAP values was calculated). For all samples CXCL9 has the highest contribution, exceeding other immunological parameters by more than twice (Figure 6(B), cyan, lime and gold bar plots). For all control samples CD40LG and PDGFB(A) follow CXCL9, with CCL22 and IL6 standing next. For ESRD samples IL6 and CSF1 are second and third, slightly outperforming PDGFB(A) (Figure 6(B), crimson bar plot). CSF1 and IL6 were previously found to be associated with kidney function and chronic kidney disease (CKD) [32, 106–108].

**Figure 6.**
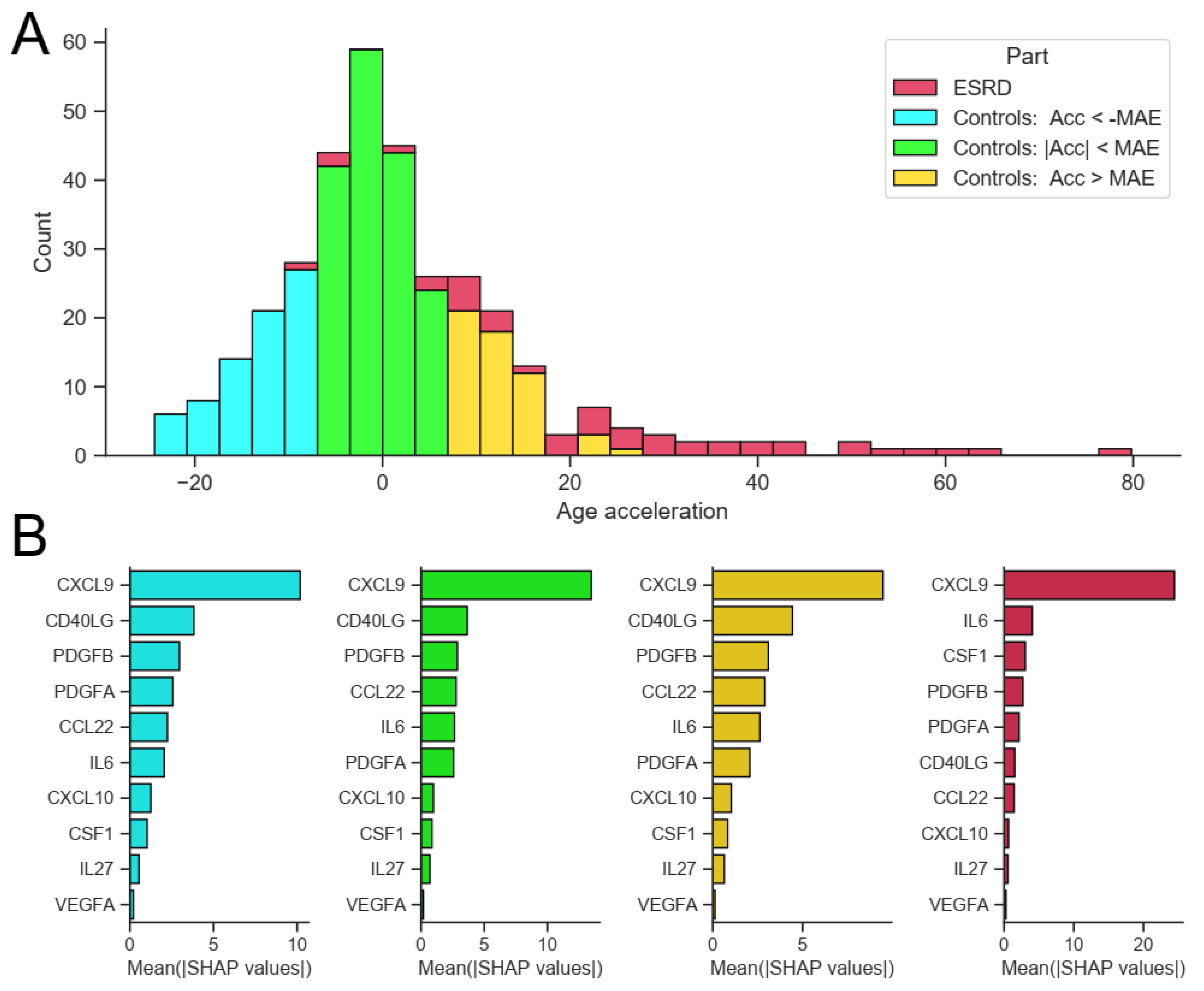
Feature importance for control samples with different types of age acceleration and ESRD samples. (A) Distribution of samples into groups: control samples with absolute value of age acceleration less than MAE (lime), control samples with value of age acceleration less than -MAE (cyan), control samples with value of age acceleration greater than MAE (gold), ESRD samples (crimson). (B) For each considered group, the bar plot illustrates the global importance of each feature, which is calculated as the average absolute value for that feature across all participating samples.

## 3. Discussion

### 3.1. Conclusion

In this paper we developed a small SImAge immunological clock that predicts a person’s age based on a limited set of immunological biomarkers. The clock shows competitive results compared to the best known high-dimensional models such as IMM-AGE [30], iAge [31], ipAge [32], and takes a smaller number of features as an input.

As a baseline model, we considered Elastic Net which is one of the most common approaches in constructing clock models based on biomedical data of different types, and also implemented various GBDT and DNN models, broadly employed in machine learning tasks on tabular data. Earlier the suitability of deep neural networks for tabular data was questioned, and it was pointed out that GBDT models often outperform them. Moreover, deep neural networks were found more difficult to optimize than GBDT models [61–63, 68]. In our study we built 12 different models with cross-validation and hyperparametric search for 46 immunological parameters, which were additionally tested on the independent dataset. Here DNN models showed better ability to generalize, outperforming GBDT models on the independent test dataset, while Elastic Net significantly underperformed both of them. As a result, the errors of the best models turned out to be lower than those of iAge and ipAGE. The next step was to reduce the model dimensionality. Since not all models are able to calculate feature importance, a unified approach based on SHAP values was used. For the best models, we ranked the features according to their summarized average SHAP values. Accordingly, 10 immunological features most robust for different models were selected. In particular, CXCL9 has been shown to play a key role in age-related chronic inflammation [31], CCL22 increases expression of pro-inflammatory mediators and decreases expression of anti-inflammatory mediators [102], IL-6 is associated with mortality risk and physical and cognitive performance [97, 109]. Based on the data of 10 selected immunological parameters, new models (4 types of models that showed the best results for the full data set - DANet, SAINT, FT-Transformer and TabNet) were built and evaluated. The FT-Transformer model showed the smallest MAE with a result of 6.94 years and Pearson *ρ* = 0.939, which is even smaller than the result for the complete data. This can be explained by filtering out the least significant noisy features, leaving only the most significant ones for age prediction. The proposed model shows significant positive age acceleration in the ESRD group in comparison to healthy controls, which may be evidence of an increased mortality risk. Local explainability methods based on SHAP values can be applied to the resulting model to obtain an individual trajectory for predicting a certain age value for each individual participant, both control and ESRD. As a result, abnormal values of certain parameters can be observed, particularly those contributing to the increase in predicted age, which may indicate the need for an in-depth medical examination. Also, subgroups of participants with different types of age acceleration were analyzed separately, and it was shown that CXCL9 has the highest contribution in all control and ESRD groups. Previously, its contribution to accelerated cardiovascular aging [31], age acceleration in Chagas disease patients [81] was shown.

Thus, we proposed an approach to build a small model of immunological age - SImAge - using the FT-Transformer DNN model, which showed the highest result among all the tested models based on 10 immunological parameters. The obtained model shows the lowest error among the published studies on immunological profile, while taking a smaller number of features as an input. Further work may include expanding the list of models tested, expanding the analyzed data through open sources, and testing the SImAge model for participants with different diseases (including age-associated and/or immunological diseases). Small size immunological panels can also prove cost efficient in practice.

### 3.2. Limitations

The proposed analysis has several limitations at its various stages. Firstly, although the developed age predictor shows better results than in [31], the size of our dataset is significantly smaller. This could be relevant for machine learning methods, in particular, since they typically perform better on large amounts of data. Nevertheless, it is interesting that machine learning techniques, which are primarily focused on big data, perform well on relatively limited data sets, like in our case. The second limitation concerns the obtained parameters of the machine learning models, that they are only locally optimal within the proposed limited hyperparametric search. It is possible that more optimal parameters exist for certain models, which may lead to a completely different ranking of the best models. Next, there are various dimensionality reduction and feature selection algorithms, while we exploited only one of them. Finally, the field of neural network architectures is actively developing and this paper necessarily considers only a limited list of the most popular ones. At the same time it should be noted that some architectures, such as TabTransformer [110] and its modifications, are not considered in the paper, since they are focused on working with categorical features. Taking only continuous data (like our immunological biomarkers) they reduced to relatively simple MLPs, whose variations are presented in the paper.

## 4. Methods

### 4.1. Data details

All possible inconveniences and risks were explained to each participant, as well as the details of the procedure. Each participant signed an informed consent and filled out a personal data processing consent, taking into account the principle of confidentiality, which implies the accessibility of personal data only to the research group. The study was approved by the local ethical committee of Lobachevsky State University of Nizhny Novgorod. All research procedures were in accordance with the 1964 Declaration of Helsinki and its later amendments. The analysis was performed on plasma using the K3-EDTA anticoagulant, without hemolysis and lipemia. Plasma was thawed, spun (3000 rpm, 10 min) to remove debris, and 25 µl was collected in duplicate. Plasma with antibody-immobilized beads was incubated with agitation on a plate shaker overnight (16–18 h) at 2–8 °C. The Luminex® assay was run according to the manufacturer’s instructions, using a custom human cytokine 46-plex panel (EMD Millipore Corporation, HCYTA-60 K-PX48). Assay plates were measured using a Magpix (Milliplex MAP). Data acquisition and analysis were done using a standard set of programs MAGPIX®. Data quality was examined based on the following criteria: standard curve for each analyte has a 5P R2 value > 0.95. To pass assay technical quality control, the results for two controls in the kit needed to be within the 95% of CI (confidence interval) provided by the vendor for > 40 of the tested analytes. No further tests were done on samples with results out of range low (< OOR). Samples with results out of range high (> OOR) or greater than the standard curve maximum value (SC max) were not tested at higher dilutions.

### 4.2. Age estimation models

This study considers various machine learning models for solving the problem of regression of chronological age using immunological profile data. All of these models focus on tabular data, for which the features have already been extracted and there is no inherent position information, which means arbitrary column order. A peculiarity of our data representation is that all of the considered immunological features are continuous (no categorical or ordinal features).

Elastic Net is a relatively simple model and a popular choice for constructing various kinds of biological clocks on tabular data such as epigenetic [26–29], immunological [32, 42], transcriptomic [43], metabolomic [44–46], microRNA [47], and proteomic [48] data.

In the last few years, a number of papers have been published comparing the effectiveness of applying GBDT and DNN to different tabular data. However, no consensus has yet been reached: some papers suggest that the result depends on the dataset size [62], some papers conclude that GBDT for tabular data works better [61, 63], and some papers propose completely new DNN architectures that perform better than GBDT [67, 68].

The following subsections present the basic concepts of the models used to predict age from immunological data.

#### Linear Model: Elastic Net

Elastic net is an extension of linear regression that adds L1 (Lasso) and L2 (Ridge) regularization penalties to the loss function during training. Linear regression assumes a linear relationship between input immunological features (*x*_1_, *x*_2_, …, *x*_*N*_) and target variable – chronological age (*y* are ground truth values and *y*^ are predictions):

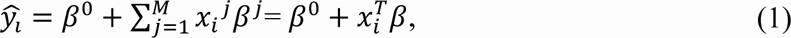

where *i* is the sample index (*i* = 1, . . ., *N*). *β*^*j*^ (*j* = 0, . . ., *M*) are coefficients in the linear model, which are found via optimization process that seeks to minimize loss function:

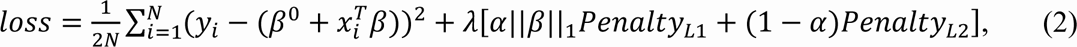

where 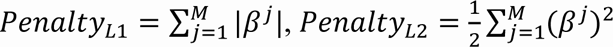, hyperparameter *λ* ≥ 0 controls the overall strength of both penalties to the loss function and 0 ≤ *α* ≤ 1 is a compromise between Ridge (*α* = 0) and Lasso (*α* = 1). In the performed experiments, equal contribution of both penalty types is set (*α* = 0.5) and only the parameter *λ* is varied.

The implementation of the algorithm was taken from the sklearn library version 1.1.2 [111]. The range of varying parameters and their precise values for the best models are presented in Supplementary Table 2 (sheet “ElasticNet”).

#### Gradient Boosted Decision Trees: XGBoost, LightGBM, and CatBoost

XGBoost (eXtreme Gradient Boosting) [58], LightGBM (Light Gradient Boosting Machine) [59], and CatBoost (Categorical Boosting) [60] are all GBDT (Gradient Boosted Decision Tree) algorithms. These models are based on ensemble learning, a technique that combines predictions from multiple models to produce predictions that are potentially more stable and better generalizable. By averaging individual model errors, the risk of overtraining is reduced while maintaining strong prediction performance. XGBoost, LightGBM, and CatBoost are variations of boosting algorithms, which build models sequentially using all data, with each model improving upon the error of the previous one. The differences between the three models lie in the splitting method and type of tree growth.

#### Splitting Method

XGBoost uses a pre-sorting algorithm which takes into account all features and sorts them by value. After that, a linear scan is performed to select the best split with the highest information gain. The histogram-based modification groups them into discrete bins and finds the split point based on these bins.

LightGBM offers gradient-based one-side sampling (GOSS) which selects the split using all instances with large gradients (higher errors) and random instances with small gradients (smaller errors). To keep data distribution, GOSS uses a constant multiplier for instances with small gradients. As a result, GOSS for learned decision trees achieves a balance between the speed of reducing the number of data points and preserving accuracy.

Catboost offers Minimal Variance Sampling (MVS) technique, which is a weighted sampling version of Stochastic Gradient Boosting (SGB) [112]. The weighted sampling occurs at tree-level, not at split-level. The observations for each boosting tree are sampled to maximize the accuracy of the split scoring.

#### Tree Growth

XGBoost splits trees up to a certain maximum depth (specified by a hyperparameter) and then starts pruning the tree backwards and removes splits beyond which there is no positive gain. It uses this approach since a split with no loss reduction can be followed by a split with loss reduction.

LightGBM uses leaf-wise tree growth: it chooses to grow a leaf that minimizes losses, allowing an unbalanced tree to grow.

Catboost builds a balanced tree: at each level of such a tree, it chooses the feature-split pair that leads to the lowest loss, and it is used for all nodes in the level.

These models also differ in imputing of missing values, processing of categorical features and computing feature importance, which is not relevant in this study: in our dataset there are no missing values and categorical features, and the feature importance is computed in a universal way for all models using SHAP values.

Software implementations of these models are used from the corresponding packages: XGBoost version 1.6.2, LightGBM version 3.3.2, CatBoost version 1.1. The range of varied parameters as well as their description and precise values for the best models are presented in Supplementary Table 2 (sheets “XGboost”, “LightGBM”, “CatBoost”). In all GBDT models, the maximum number of rounds of the training process was set to 1000, and the number of early stopping rounds was set to 100 (models stopped the training process if the evaluation metric in the test dataset was not improving for 100 rounds).

#### Deep Neural Networks

Several implementations of neural network architectures focused on tabular data are used in this work. The concepts and basic elements of the deep neural network architectures used in this work, taking into account the basic details of their software implementations, are presented in Figure 7. For training all neural network models, the Adam optimization algorithm was used [113], whose main hyperparameters (learning rate and weight decay) were selected for each model individually during the hyperparametric search. Their ranges and precise values for the best models are presented in Supplementary Table 2.

**Figure 7.**
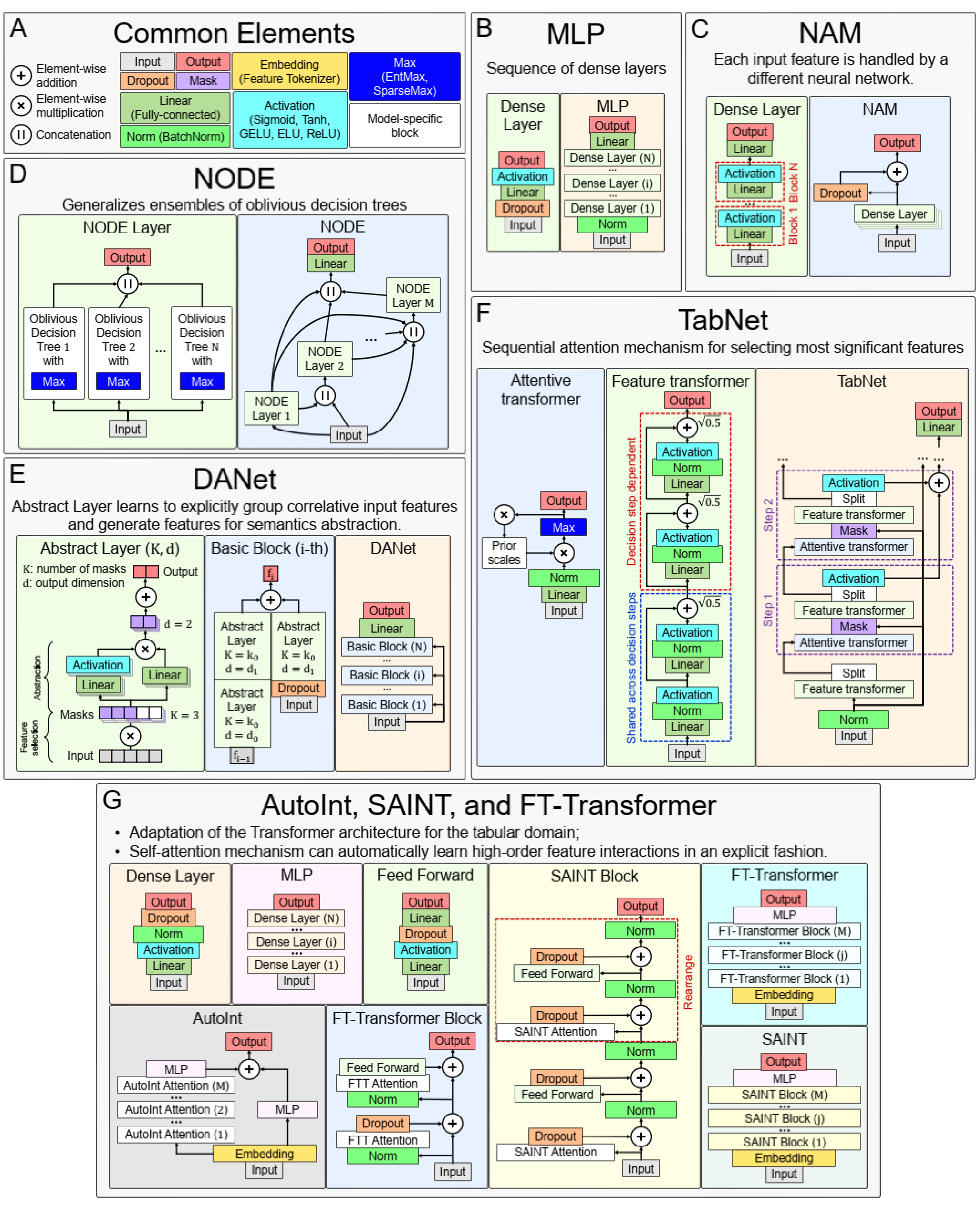
Main concepts of neural network architectures used for chronological age regression on immunological parameters. (A) Blocks and layers used in many architectures. (B) Multilayer Perceptron (MLP) - the simplest neural network architecture composed of a sequence of dense layers. (C) Neural Additive Model (NAM) - the architecture where a separate neural network is built for each input feature. (D) Neural Oblivious Decision Ensembles (NODE) - the architecture that combines ensembles of oblivious decision trees with the benefits of both end-to-end gradient-based optimization and the power of multi-layer hierarchical representation learning. (E) Deep Abstract Network (DANet), based on special abstract layers that learn to explicitly group correlative input features and generate features for semantics abstraction. (F) TabNet architecture, which implements a sequential attention mechanism to select the most important features. (G) AutoInt, SAINT, and FT-Transformer are conceptually similar neural network architectures, based on adaptation of Transformer architecture for the tabular domain.

For all models, the maximum number of epochs has been set to 1000, and for early stopping the number of epochs has been set to 100 (models stopped the training process if the evaluation metric on the test dataset was not improving for 100 epochs). All models were implemented in the PyTorch Lightning framework [114], which is a lightweight PyTorch [115] wrapper for high-performance AI research. Feature importance for all models were calculated using SHAP values and corresponding global explainability methods [95] (see Section 4.4 for details).

Common Elements.

Element-wise addiction, multiplication and concatenation (of output vectors) are used in many neural network architectures to variously accumulate outputs of several previous layers at once. Dropout is a special layer that provides regularization and prevents co-adaptation of neurons by zeroing out randomly selected neurons [116]. It works only during training and is turned off in evaluation mode.

Mask layers are used for instance-wise feature selection in several considered models (TabNet and DANet).

Linear (or fully-connected, or dense) layer applies a linear transformation to the input data with an optional bias. It is often used in the final stages of neural networks (but not only in them) to change the dimensionality of the output of the preceding layer, so that the model can easily determine the relationship between the data values in which the model operates.

Normalization layers bring all input values to the same scale with a mean equal to zero and variance equal to one. This improves productivity and stabilizes neural networks.

Embedding layers transform input continuous features into a new embedding space of a different dimensionality, using both linear and nonlinear transformations. During the neural network training, similar samples can be clustered in this space.

Activation functions are used to activate neurons from hidden layers and to introduce different nonlinearities in the decision boundary of the neural networks. In many used models the type of activation function is a hyperparameter.

EntMax [117] and SparseMax [118] are modifications of activation functions that transform continuous vectors into a discrete probability distribution, resulting in a sparse output. They are used to implement the attention mechanism in some models, allowing to take into account the influence of only the most important features.

The designations of the described general blocks in the general scheme of the neural network concepts used in this work are shown in Figure 7(A).

Multilayer Perceptron (MLP).

A simple neural network architecture, consisting of several dense blocks, which in turn consist of linear, dropout, and activation layers, as shown in Figure 7(B).

The software implementation of the neural network is adapted from the pytorch-widedeep library version 1.2.1. The main hyperparameters are the architecture of dense blocks, activation function type, and probability of dropout. Exact values of the varied parameters are presented in the Supplementary Table 2 (sheet “MLP”).

Neural Additive Model (NAM).

In this architecture [93], a separate dense block consisting of linear layers and activation functions is constructed for each input feature. All these independent dense blocks are trained jointly and can learn arbitrarily complex relations between their input and output features. The main advantage of this method is its ease of interpretability, as each input function is processed independently by a different neural network. A scheme of the architecture is shown in Figure 7(C).

#### The software implementation of the neural network is adapted from the NAM library version

0.0.3 implementing this architecture for the PyTorch framework. The hyperparameters of the network are the architecture of dense blocks, activation function type, probability of dropout and additional regularization parameters [93]. Exact values of the varying parameters are given in the Supplementary Table 2 (sheet “NAM”).

Neural Oblivious Decision Ensembles (NODE).

This architecture [69] combines decision trees and deep neural networks, so that they can be trained (via gradient-based optimization) in an end-to-end manner. This method is based on so-called oblivious decision trees (ODTs), a special type of decision tree that uses the same splitting function and splitting threshold in all internal nodes of the same depth (as in CatBoost [60]). It uses EntMax transformation [117], which effectively performs soft splitting feature selection in decision trees within the NODE architecture. A scheme of the architecture is shown in Figure 7(D).

The software implementation of the neural network is adapted from the PyTorch Tabular package version 0.7.0 [119]. The hyperparameters of the network are number of NODE layers, numbers of ODTs in each NODE layer, the depth of ODTs, sparse activation function type. Exact values of varying parameters are presented in Supplementary Table 2 (sheet “NODE”). Deep Abstract Network (DANet).

The architecture [94] is focused around abstract layers, the main idea of which is to group correlated features (through a sparse learnable mask) and create higher level abstract features from them. The DANet architecture consists of stucking such abstract layers into blocks. The blocks are combined one by one, and each block has a shortcut connection that adds raw features back to each block. A scheme of the architecture is shown in Figure 7(E).

The software implementation of the neural network is adapted from the supplementary code repository of the corresponding article [94]. The hyperparameters of the network are number of abstract layers to stack, number of masks, the output feature dimension in the abstract layer, and dropout rate in the shortcut module. Exact values of the varying parameters are presented in the Supplementary Table 2 (sheet “DANet”).

TabNet.

The architecture [65] consists of sequential modules (steps), each implementing a sequential attention mechanism that selects the most significant features. Attentive transformers learn the relationship between the relevant features and decide which features to pass using the SparseMax (or EntMax) functions. In the feature transformer block, all the selected features are processed to generate the final output. Each feature transformer is composed of several normalization layers, linear layers and several Gated Linear Unit (GLU) blocks [120], that control which information should pass through the network. The scheme of the architecture is shown in Figure 7(F).

The software implementation of the neural network is adapted from the pytorch-widedeep library version 1.2.1. The hyperparameters of the network are width of the decision prediction layer, number of decision steps, number of GLU blocks, parameters of batch normalization, and relaxation parameters. Exact values of the varying parameters are presented in Supplementary Table 2 (sheet “TabNet”).

AutoInt.

This model tries to automatically learn the interactions between features and create a better representation, and then use that representation in downstream tasks. The model first transforms the features into embeddings, and then applies a series of attention-based transformations to the embeddings. The output of the model is the sum of the outputs of the multi-head self-attention mechanism and the residual connection block. A scheme of the architecture is shown in Figure 7(G).

The software implementation of the neural network is adapted from the PyTorch Tabular package version 0.7.0 [119]. The hyperparameters of the network are different characteristics of multi-headed attention layers, dropout rates, and linear layers configuration in MLP. Exact values of the varied parameters are presented in the Supplementary Table 2 (sheet “AutoInt”). Self-Attention and Intersample Attention Transformer (SAINT).

This hybrid architecture is based on self-attention, which applies attention to both rows and columns, and includes an enhanced embedding method [67]. SAINT projects all features into a combined dense vector space. These projected values are passed as tokens into the transformer encoder, which performs a special attention mechanism. A scheme of the architecture is shown in Figure 7(G).

The software implementation of the neural network is adapted from the pytorch-widedeep library version 1.2.1 and uses Einstein notation for operations on tensors (rearrange from einops python package [121]). The hyperparameters of the network are the number of attention heads per transformer block, the number of SAINT-transformer blocks, dropout rates, and linear layers configuration in MLP. Exact values of the varying parameters are presented in Supplementary Table 2 (sheet “SAINT”).

Feature Tokenizer and Transformer (FT-Transformer).

Like the previous two models, it is an adaptation of the transformer architecture [122] for the tabular domain. First, in this architecture, the function tokenizer transforms features into embeddings. Then the embeddings are processed by the Transformer layer stack with special multi-head self-attention blocks. A scheme of the architecture is shown in Figure 7(G).

The software implementation of the neural network is adapted from the pytorch-widedeep library version 1.2.1. The hyperparameters of the network are number of attention heads per transformer block, number of FT-Transformer blocks, dropout rates, and linear layers configuration in MLP. Exact values of the varying parameters are presented in Supplementary Table 2 (sheet “FT-Transformer”).

### 4.3. Experiment details

#### Cross-validation and metrics

Cross-validation is a resampling procedure used to evaluate machine learning models on a limited amount of data. In this paper, we used a k-fold cross-validation procedure with k=5. The mean result over several splits is expected to be a more accurate estimate of the true unknown underlying mean performance of the model in the dataset, calculated using standard error.

The software implementation of the k-fold cross-validation procedure was used from the sklearn library version 1.1.2 [111]. Stratification was performed as follows: the whole age range was divided into four bins of equal length and the samples inside each bin were divided into almost equal five splits.

Various metrics such as MAE [32, 43, 45, 46, 52, 53, 53–56], RMSE [26, 45], correlation between chronological age and predicted age [28, 46, 48], median error [27, 51], and others are used to evaluate the efficiency of machine learning models in solving age estimation problems. The most popular among them is MAE. In this work, MAE was also chosen as the main observable metric. It is calculated by the formula:

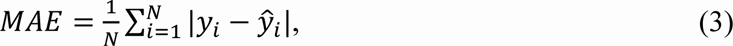

where *y*_*i*_ – ground truth values of target metric (chronological age), *y*^_*i*_ – model predictions, *N* – number of samples. For this metric two values are given for each model: (i) the main metric by which models are sorted – average MAE (±STD) for all cross-validation splits; (ii) the secondary metric – MAE for the best model (for one particular split, where the best results were obtained).

Pearson correlation coefficient was also tracked along with the MAE. This helps to avoid incorrect model performance, when it predicts age with a low error only for the most representative age range, and for all others, the error is high. In such cases, the correlation coefficient is low. Pearson correlation coefficient is calculated by the formula:

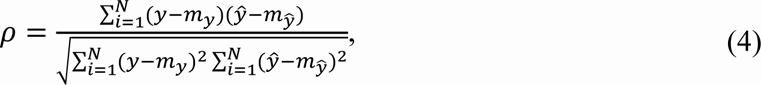

where *y* – ground truth values of target metric (chronological age), *y*^ – model predictions, *N* – mean of the vector *y*, *m*_*y*_ – mean of the vector *y*^. For each model, average *ρ* (±STD) for all cross-validation splits as well as *ρ* for the best split is given.

First, the original dataset was divided into training and validation ones in an 80/20 ratio, and a cross-validation procedure with hyperparametric search was performed to identify the best model (with the lowest MAE) with optimal parameters for each model type. The selected models were then tested on the independent dataset, and the best MAE was selected among them on the test dataset. In both cases, the value of the Pearson correlation coefficient was also tracked for the best models.

#### Hyperparameters optimization

Each machine learning model considered in this paper has certain hyperparameters. To obtain the best result, hyperparameter search of the best combination was performed, which yielded the minimum MAE value averaged over all the cross-validation splits.

The software implementation of hyperparameter optimization was taken from the optuna python package version 3.0.2 [123], using the Tree-structured Parzen Estimator (TPE) algorithm [124]. TPE is an iterative process that uses the history of evaluated hyperparameters to create a probabilistic model that is used to propose the next set of hyperparameters for evaluation. The total number of optimization trials for each considered model was set to 200. The number of random sampled startup trials was set to 50. The number of candidate samples used to calculate the expected improvement was set to 10. Supplementary Table 2 lists the varied hyperparameters for each model with a description and range of variation.

### 4.4. SHAP values

SHAP (Shapley Additive ExPlanations) values are a game theory approach to explain model predictions. SHAP values for a regression problem have the same dimensionality as the original dataset - they are computed for each sample and for each feature. SHAP values show how a particular value of a selected feature for a particular sample changed the baseline model prediction (the average model prediction on the background dataset - train and validation datasets together in our case).

The model output can be interpreted as the reward that will be distributed among the group of features that helped obtain the reward. SHAP values determine how the model output, the reward, should be distributed among the features. Given the prediction of model *f* on sample *x*_*i*_ the method estimates SHAP values ϕ(*x*_*i*_, *f*) by finding an approximation to the model, *g*_*i*_(*x*_*i*_), one model per sample *x*_*i*_. This model *g*_*i*_(*x*_*i*_) is locally accurate: its output converges to *f*(*x*_*i*_) for this sample by summing up *M* attributions ϕ_*j*_(*x*_*i*_, *f*) for *M*features. The model *g*_*i*_(*x*_*i*_) is consistent: features that are truly important to one model’s predictions versus another are always assigned higher importance. The summarization of effects:

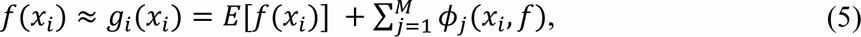

where *E*[*f*(*x*_*i*_)] is the expected reward across the training samples [125]. SHAP values can be computed as a weighted sum of the contribution of a feature to the model prediction, taking into account all possible permutations of other features introduced into the model. This method requires significant computational resources, so various estimators of SHAP values (explainers) are implemented in the shap python package version 0.41.0 [95].

For GBDT and DNN models, [95] presents special types of explainers, TreeExplainer and DeepExplainer, respectively. DeepExplainer does not support specific layers of some architectures (for example, SparseMax and EntMax in TabNet and Node). For this reason, the KernelExplainer is used as unified as possible for all model types.

## Data availability statement

All data generated during this study are included in this published article and its supplementary files.

## Code availability statement

The source code for the analysis pipeline presented in the manuscript is publicly available.

Project name: SImAge

Project home page: https://github.com/GillianGrayson/SImAge

Operating system(s): Platform independent

Programming language: Python

Other requirements: Python 3.9, torch 1.10, pytorch-lightning 1.6.4, xgboost 1.7.0, catboost 1.1.1, lightgbm 3.3.3, pytorch-tabnet 3.1.1, scikit-learn 1.1.3, pytorch-widedeep 1.1.1, shap 0.39.0. All requirements are listed in the requirements.txt file in the project home page. License: MIT

## Abbreviations

AI: Artificial Intelligence
AutoInt: Automatic Feature Interaction Learning via Self-Attentive Neural Network
CatBoost: Categorical Boosting
CI: Confidence Interval
CKD: Chronic Kidney Disease
DANet: Deep Abstract Network
DNA: Deoxyribonucleic Acid
DNN: Deep Neural Network
ESRD: End-Stage Renal Disease
FDR: False Discovery Rate
FT-Transformer: Feature Tokenizer and Transformer
GBDT: Gradient-Boosted Decision Tree
GLU: Gated Linear Unit
GOSS: Gradient-based One-Side Sampling
LightGBM: Light Gradient Boosting Machine
MAE: Mean Absolute Error
MLP: Multilayer Perceptron
MVS: Minimal Variance Sampling
NAM: Neural Additive Model
NODE: Neural Oblivious Decision Ensemble
ODT: Oblivious Decision Tree
OOR: Out Of Range
RMSE: Root Mean Squared Error
SAINT: Self-Attention and Intersample Attention Transformer
SGB: Stochastic Gradient Boosting
SHAP: Shapley Additive Explanations
STD: Standard Deviation
TPE: Tree-structured Parzen Estimator
XAI: Explainable Artificial Intelligence
XGBoost: eXtreme Gradient Boosting

## Author’s Contributions

Conceptualization: A.K., I.Y., M.V., M.I.; Investigation: E.K.; Formal analysis: A.K., I.Y.; Methodology: A.K., I.Y., M.I.; Software: A.K., I.Y.; Resources: E.K., M.V.; Supervision: M.G.B., C.F., M.V., M.I.; Visualization: A.K., I.Y.; Writing – original draft: A.K., I.Y.; Writing – review & editing: A.K., I.Y., E.K., M.G.B., C.F., M.V., M.I.

## Acknowledgements

The authors acknowledge the use of computational resources provided by the “Lobachevsky” supercomputer. We thank the members of the Branch FESFARM NN headed by Nadezhda Lobanova for accompanying patients on hemodialysis.

